# Design of 18 nm Doxorubicin-loaded 3-helix Micelles: Cellular Uptake and Cytotoxicity in Patient-derived GBM6 Cells

**DOI:** 10.1101/2020.05.04.072892

**Authors:** Benson T. Jung, Katherine Jung, Marc Lim, Michael Li, Raquel Santos, Tomoko Ozawa, Ting Xu

## Abstract

The fate of nanocarrier materials at the cellular level constitutes a critical checkpoint in the development of effective nanomedicines, determining whether tissue level accumulation results in therapeutic benefit. The cytotoxicity and cell internalization of ~18 nm 3-helix micelle (3HM) loaded with doxorubicin (DOX) was analyzed in patient-derived glioblastoma (GBM) cells *in vitro*. The inhibitory concentration (IC_50_) of 3HM-DOX increased to 6.2 µg/mL from < 0.5 µg/mL for free DOX in patient derived GBM6 cells, 15.0 µg/mL from 6.5 µg/mL in U87MG cells, and 21.5 µg/mL from ~0.5 µg/mL in LN229 cells. Modeling analysis of previous 3HM biodistribution results predict these cytotoxic concentrations are achievable with intravenous injection in rodent GBM models. 3HM-DOX formulations were internalized intact and underwent intracellular trafficking distinct from free DOX. 3HM was quantified to have an internalization half-life of 12.6 h in GBM6 cells, significantly longer than comparable reported liposome and polymer systems. 3HM was found to traffic through active endocytic processes, with clathrin-mediated endocytosis being the most involved of the pathways studied. Inhibition studies suggest substantial involvement of low density lipoprotein receptor (LDLR) in initiating 3HM uptake. Since 3HM surface is polyethylene glycol (PEG)-ylated with no targeting functionalities, protein corona involvement in 3HM recognition is expected. The present work develops insights of the cytotoxicity, pharmacodynamics and cellular interactions of 3HM and 3HM-DOX relevant for ongoing pre-clinical studies. This work also contributes to efforts to develop predictive mathematical models tracking accumulation and biodistribution kinetics at a systemic level.

**Graphical Abstract:** 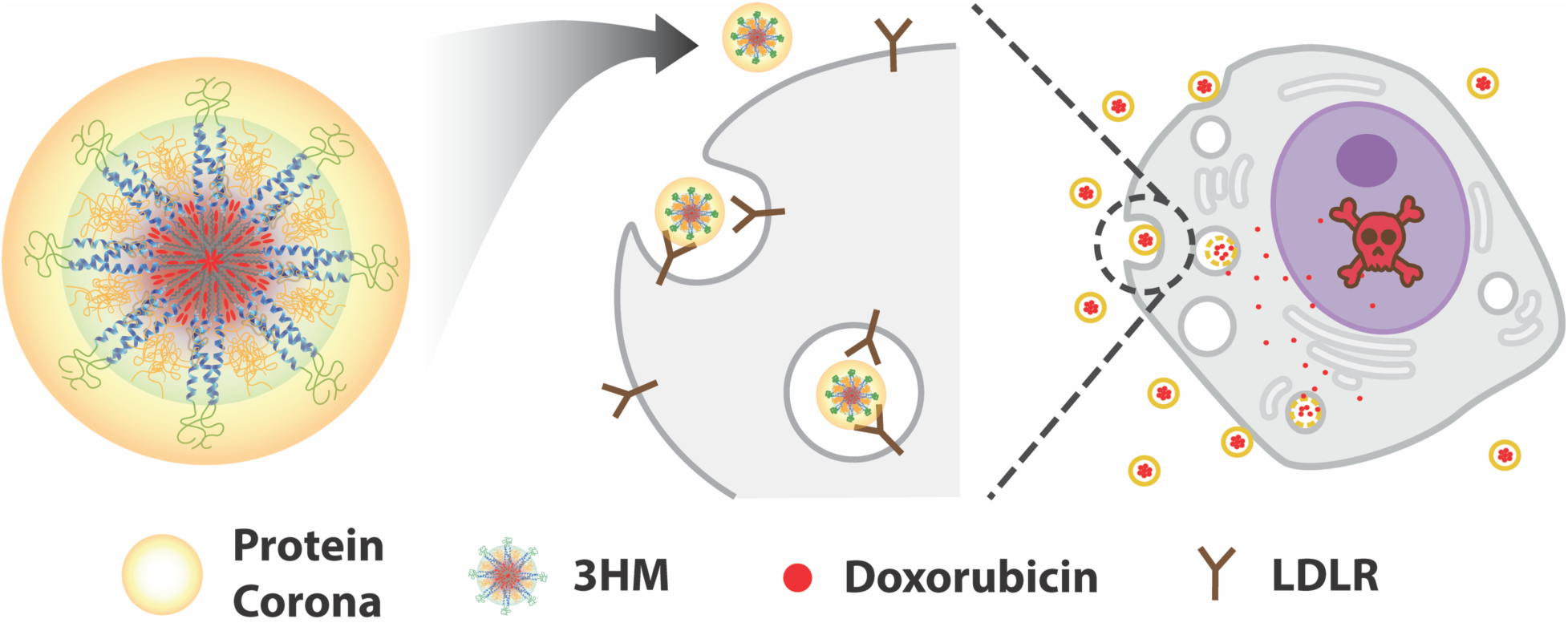

## 1. Introduction

Nanocarriers, in particular those less than 50 nm in size, are attractive delivery systems to increase the accumulation of drugs in tumors, and decrease systemic off-target delivery. ^1–5^ Peptide-polymer conjugate amphiphiles form uniform ~18 nm highly stable micelles, called 3-helix micelles (3HM). ^6–10^ 3HM has been shown to cross the disrupted blood brain barrier (BBB) from intravenous injection in rodent models and penetrate deeply into U87MG 10,11 and GBM6 orthotopic glioblastoma (GBM) tumors. ^6–8,10,12^ *In vivo* studies have demonstrated 3HM has long blood circulation, ^7,10^ significantly reduced uptake in off-target organs, ^7,10^ and lowered toxicity when formulated with doxorubicin (DOX). ^8^ Detailed characterization of the internal structure and dynamics of 3HM have laid the groundwork to understand the promising biodistribution of 3HM, with small size, and high kinetic stability being key to its performance. ^6,13–16^ However, the interactions of 3HM with target GBM cells have yet to be elucidated. Can 3HM formulations deliver active drug cargo once reaching tumor cells? How do GBM (or any) cells manipulate 3HM, if at all? What role do 3HM’s physical design features play in controlling its cellular fate? *In vitro* studies allow these questions to be answered, bridging the gap between the molecular architecture and *in vivo* performance of 3HM.

Pursuing 3HM for GBM treatment is further motivated by the clinical reality of the disease: median survival remains ~15 months even with treatment. ^17^ Multi-level obstacles to transport, including crossing the BBB, prevent most chemotherapeutics from even accessing GBM tumors. ^18,19^ At the cellular level, the defining challenge of GBM is its heterogeneity across patients, initial genetic mutations, and microenvironmental niches. ^17,18^ *In vitro*, this heterogeneity manifests in two main issues: differences in drug transport to its intracellular molecular target and cell-specific susceptibility, which can still be tied to transport variability between cell populations. ^17,20^ The present study uses patient-derived xenograft (PDX) cell models *in vitro* to capture as much of the variability in transport and susceptibility of GBM cells that occurs in the clinical case as possible. ^17,21^ PDX cells were harvested from flank tumors and subsequently cultured in 2D for short passage numbers, where they maintain their pathological heterogeneity (dense, angiogenic, and invasive niches) once re-implanted orthotopically in mice, as shown by Garner et al. ^22^

The PEGylation architecture of 3HM is composed of two chains of different molecular weights and conjugation sites, and may afford behavior in biological contexts different from other homogeneously PEGylated nanocarriers. ^6,13,14^ As shown in Scheme 1, there is a 0.8 nm brush layer of PEG (MW:750 g/mol, ‘PEG750’) attached to the end of 3-helix bundle. ^10^ Side-conjugated PEG (MW:2000 g/mol, ‘PEG2K’) is presented laterally in a regular geometric arrangement by the peptide coiled-coil structure. MD simulations and structural scattering studies published previously suggest the PEG2K are exposed on the surface of the micelles in between the PEG750 brushes. ^6,10,13^ The resulting patterned surface of 3HM may be pivotal in mediating its interaction with media/serum biomolecules and subsequent biological machinery.

**Scheme 1:**
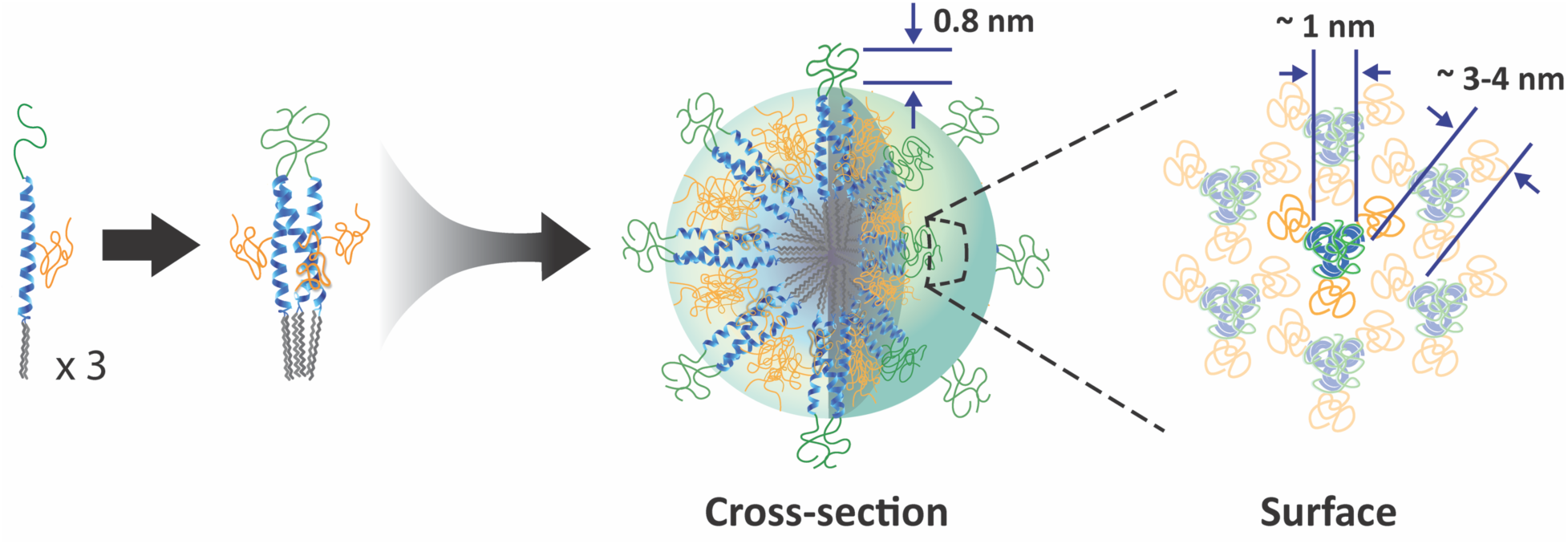
Illustration of the lateral correlation of PEG750 and PEG2K position with peptide architecture in 3HM. Peptide is shown in blue. Alkyl tails are shown in gray. PEG750 is shown in green. PEG2K is shown in yellow.

The current work explores the cellular fate of 3HM formulations, elucidating their cell uptake mechanisms, kinetics, and cytotoxicity in the context of GBM therapy development. Investigation of 3HM interactions with specific cellular machinery once introduced into biological contexts provides insights for future nanocarrier design efforts. Investigation of the kinetics and molecular mechanisms of particle uptake supports and informs ongoing *in vivo* preclinical work. This work also contributes to efforts to develop predictive mathematical models tracking accumulation and biodistribution kinetics at a systemic level. ^23^

## 2. Experimental Methods

### 2.1. Synthesis and labeling of 3HM Peptide−Polymer Amphiphiles

The synthetic and purification procedures of the 3HM amphiphile have been previously documented in detail. ^7,12^ Briefly, the 3-helix bundle forming peptide, 1CW (EVEALEKKVAALEC KVQALEKKVEALEHGW) was modified at the N-terminus with Fmoc-6-aminohexanoic acid (Ahx) as a linker, followed by a Fmoc-Lys(Fmoc)-OH residue, and at the C-terminus with additional residues Fmoc-Gly-Gly-Gly-Lys(Alloc)-OH. N-terminal modification allowed for conjugation of two alkyl tails (C18) leading to the amphiphilic molecule. The cysteine residue at position 14 facilitated conjugation of a maleimide-PEG2K to the amphiphile that provides entropic stabilization previously reported in detail. ^6^ The modified peptide (K(Fmoc)-Ahx-EVEALEKKVAALECKVQALEKKVEALEHGWGGGK(Alloc)) was produced on a Prelude solid phase peptide synthesizer (Protein Technologies, AZ) using 9-fluorenylmethyl carbamate (Fmoc) chemistry. The N-terminal alkyl chains were conjugated through reaction of stearic acid (C18) with deprotected Fmoc-Lys(Fmoc)-OH. The C-terminal Fmoc-Lys(Alloc)-OH was selectively deprotected with five 30 min reactions of 0.2 equiv. of Pd(PPh_3_)_4_ used as a catalyst, with 24 equivalents of phenylsilane as an allyl acceptor in dichloromethane. The resulting free amino group was utilized for conjugating carboxy-terminated PEG750 or 5,6-carboxyfluorescein (FAM) (as a fluorescent tag) using HBTU/DIPEA chemistry. Cleavage was carried out using a cocktail of 95:2.5:2.5 TFA/TIS/water for 3 h. Crude peptides were precipitated and washed three times in cold ether, isolated, and dried. Maleimide-Cys conjugation was used to specifically conjugate PEG2K to the cysteine at position 14 of the peptide sequence. The conjugation reaction was carried out in 9:1 HEPES buffer (100 mM, pH=7.4):MeOH overnight under nitrogen with a reaction ratio of PEG to peptide at 2:1. The final peptide amphiphiles (MW= ~7200 g/mol) were isolated with centrifugal filters (Amicon, MW cutoff: 3000 g/mol). Spin filtration was performed at 7000 rpm for 40 min, and the concentrate was washed with deionized water. This spin procedure was repeated seven times, then the amphiphiles were lyophilized to a powder. Peptide amphiphile masses were confirmed with Matrix-Assisted Laser Desorption Ionization Time of Flight (MALDI-TOF) (Supporting Information, Figure 1).

**Figure 1.**
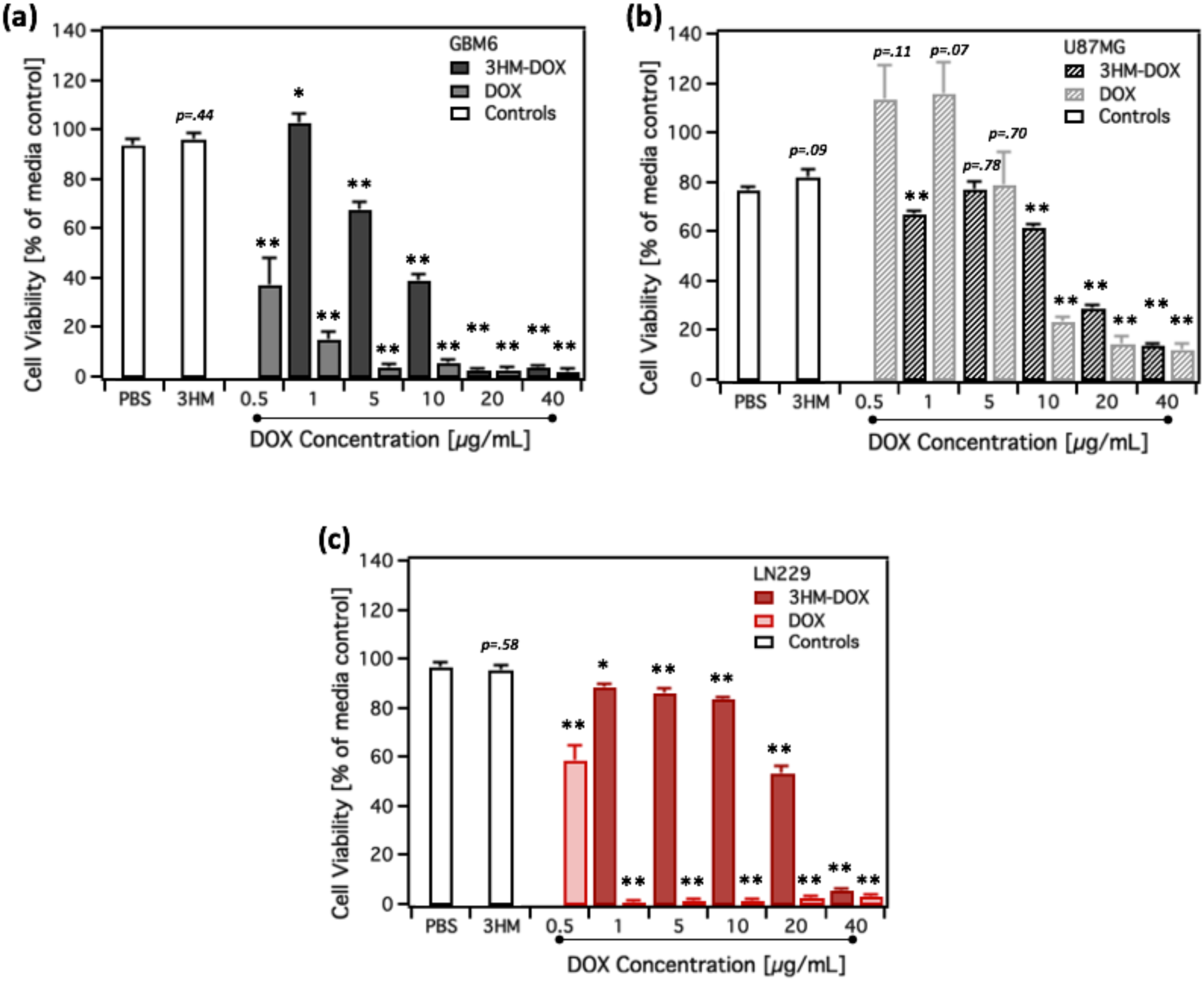
(a) GBM6, (b) U87MG (c) LN229 viability after 48 h treatment with free DOX and 3HM-DOX followed by a 72 h recovery period after treatment removal before assaying by MTT. Blank 3HM (62.5 µg/mL) were concentration matched to 5 µg/mL 3HM-DOX samples. * *p* <. 05, ** *p* <. 001 for series vs respective PBS controls.

### 2.2. Doxorubicin Encapsulation

The thin film hydration method to encapsulate 3HM with DOX has been described previously. ^8^ Briefly, purified 3HM was dissolved in methanol to 20 mg/mL. 0.2 mg/mL DOX in methanol was added to 13 wt% of 3HM. The co-dissolved 3HM and DOX was sonicated and dried to a film under airflow in darkness at room temperature. Residual methanol was evaporated in a vacuum oven at 60 °C for 2 h. Phosphate buffer (25 mM, pH 7.4) was used to rehydrate the film to ~1 mg/mL. The micelle solution was annealed at 70 °C for 45 min and stirred at room temperature overnight to equilibrate the drug loaded micelles. Unencapsulated DOX was removed with centrifugal filters (Amicon, MW cutoff: 3000 g/mol). Spin filtration was performed at 7500 rpm for 30 min, and the retentate was washed with deionized water. This step was repeated until the DOX loading, as measured by UV-vis absorbance (wavelength: 480 nm), reached a constant value (~8-9 wt%) indicating all free drug had been removed. The retained 3HM-DOX concentrate was lyophilized for storage. For further experiments, 3HM-DOX was dissolved in phosphate buffer (25 mM, pH 7.4) to the desired concentration, annealed at 70 °C for 45 min, and allowed to cool to room temperature before use.

### 2.3. Differential Scanning Calorimetry

DSC Thermograms were obtained for buffered 3HM and 3HM-FAM solutions (~1 mg/mL in phosphate buffer, 25 mM, pH 7.4) using a VP-Microcal calorimeter (GE). ~550 µL samples were loaded and equilibrated at 5 °C for 15 min. Temperature was increased to 55 °C at a scan rate of 1 °C/min. Thermograms were baseline corrected, normalized for exact concentration, and fit using Origin software provided with the VP-Microcal instrument.

### 2.4. Fluorescence Recovery

Fluorescence spectroscopy was performed on 3HM-FAM solutions using a LS-55 fluorescence spectrometer (PerkinElmer). Stock solutions of FAM-labeled amphiphiles and unlabeled 3HM were prepared to 1 mg/mL in phosphate buffer (25 mM, pH 7.4). The two solutions were mixed to give a final molar ratio of 2:43 FAM: unlabeled 3HM. The mixed solution was diluted to 0.2 mg/mL and annealed at 70 °C for 45 min. After annealing, the micelle solution was equilibrated at room temperature for 1 h to homogenize FAM-labeled amphiphile distribution through the micelles before fluorescence recovery experiments. 3HM-FAM solutions of the same molar ratio of labeled/unlabeled amphiphiles were mixed with phenol free DMEM (Gibco, 4.5 g/L D-glucose, 10 % FBS, 1% L-glutamine, 1% Sodium pyruvate, 1% NEAA, 1% amphotericin B, 1% penicillin−streptomycin) at a ratio of 1:3 to mimic subsequent cell studies. Samples were loaded into a 1 mm path length quartz cell (Starna Cells). Emission spectra were recorded from 490 nm to 640 nm using an excitation wavelength of 450 nm at a scan rate of 200 nm/min. Temporal evolution of fluorescence intensity was recorded every 15 min at 522 nm. Temperature was controlled with a Peltier temperature controller (PTP-1, PerkinElmer). Fluorescence recovery experiments were conducted with the sample cell maintained at 37 °C for 48 h. Fluorescence intensity was normalized to initial fluorescence at time zero.

### 2.5. Cell Culture

Human GBM primary tissue, GBM6, were maintained as serially passaged subcutaneous xenografts in athymic mice in the Preclinical Therapeutic Testing Core Facility at the Brain Tumor Center at the University of California, San Francisco. ^21^ Harvested GBM6 flank tumors were finely minced and incubated with papain for 30 min to produce a tissue suspension. This suspension was subsequently filtered through 70 µm and 45 µm strainers. The filtered cell suspension was centrifuged at 1000 rpm for 10 min to produce a cell pellet. Supernatant was removed, and cells were resuspended in D-PBS (-/-). The cell suspension was again centrifuged at 1000 rpm for 10 min. Supernatant was removed and replaced with 1 mL fresh media. Cells were plated in phenol free DMEM (Gibco, 4.5 g/L D-glucose, 10 % FBS, 1% L-glutamine, 1% Sodium pyruvate, 1% NEAA, 1% amphotericin B, 1% penicillin−streptomycin) and incubated for 2 days (37 °C in 5% CO_2_). Media was changed to remove nonadherent cells and residual debris. Cells were passaged at 80% confluency 5 times before being used for experiments to remove non-target primary cells. Cells were not used past 20 passages. LN229 and U87MG immortalized GBM cells (Cell Culture Facility, University of California, Berkeley) were cultured with similar protocols.

### 2.6. *In vitro* metabolic activity by MTT assay

MTT (3-(4,5-dimethylthiazol-2-yl)-2,5-diphenyltetrazolium bromide) assay was used to study the viability of cells treated with 3HM formulations. Cells were seeded in triplicate in 96 well plates at 2000 cells/well (culture volume 100 µL). GBM6 cells were incubated for 48 h before treatment to reach similar confluency (70%) as the other cells (LN229, U87MG), which were incubated for 24 h. Cell media was removed, and treatments (media, phosphate buffer, blank 3HM, 3HM-FAM, DOX-loaded 3HM or free DOX) were introduced at a 1:3 ratio in media. 3HM and 3HM-FAM were concentration matched to the peptide content of the DOX-loaded micelles. Treatments were incubated for 48 h, then removed, replaced with fresh media, and incubated an additional 72 h. Viability of cells exposed to suramin, used for uptake pathway determination, was done with the same procedure. MTT reagent solution (1.2 mM in PBS (+/+), 100 µL) was added to the wells and incubated for 2 h. Media and MTT solution was removed, and 100 µL of DMSO was added to each well and incubated (37 °C) for 10 min to dissolve formazan crystals homogeneously. Absorbance was measured at 570 nm using an Infinite M200 microplate reader (Tecan). Absorbance values were normalized to media-only controls to represent cell viability.

### 2.7. Confocal Fluorescence Microscopy

Confocal microscopy was performed using a Zeiss LSM 880 NLO AxioExaminer (CRL Molecular Imaging Center, University of California, Berkeley). GBM6 cells were seeded on 8-well glass culture slides and allowed to reach the desired confluency with culture conditions above. Cells were incubated with 3HM-DOX (final DOX conc. 5 µg/mL) and 3HM-FAM (final conc. ~62.5 µg/mL) in a ratio of 1:3 in media. If used, Lysotracker dye (Invitrogen, Carlsbad, CA) was incubated with cells for 30 min. Cells were washed in DPBS (-/-) 3 times before being fixed with 4% paraformaldehyde for 10 min. Cells were washed 3 times with 0.1 M glycine in PBS (-/-) followed by PBS (-/-). DNA was stained with 1 X DAPI (900 nM) for 5 min and washed 3 times with PBS (-/-). Slides were mounted with Fluoromount-G and imaged within the same day.

### 2.8. Nanoparticle Internalization by Flow Cytometry

Flow cytometry (FCT) was used to measure the internalization of fluorescently labeled 3HM. Cells were seeded in triplicate in 12 well plates at 10000 cells/well (culture volume 1 mL). Cells were incubated for 48 h to allow attachment. Cell media was removed, and treatments were introduced at a 1:3 ratio in media: media, phosphate buffer, blank 3HM, 3HM-FAM, and DOX-loaded 3HM. 3HM and 3HM-FAM were concentration matched to the peptide content of the DOX-loaded micelles. Treatments were continuously incubated with cells for 1 min - 48 h. Cells were washed with PBS (-/-) and harvested with 1X Accutase (Stemcell Technologies) incubation at 37 °C for 10 min. Media was added, and the cell suspension was centrifuged at 1000 rpm for 5 min. Supernatant was discarded, and cells were resuspended in cold PBS (-/-) with 2.5% FBS. Cells were strained, counted, and diluted, if necessary, to ~1×10^7^ cells/mL. FCT samples were kept on ice and measured immediately after preparation. An LSR II Analyzer (BD Biosciences) was used to measure 10,000 cells for each well. FAM fluorescence was measured at 530 nm, and DOX fluorescence was measured at 575 nm, with an excitation wavelength of 488 nm for both channels. Cell gating for singlet viable cells was performed and verified with propidium iodide staining (Supporting Information, Figure 2). Average bulk fluorescence from each gated sample was background-reduced using the un-treated samples.

**Figure 2.**
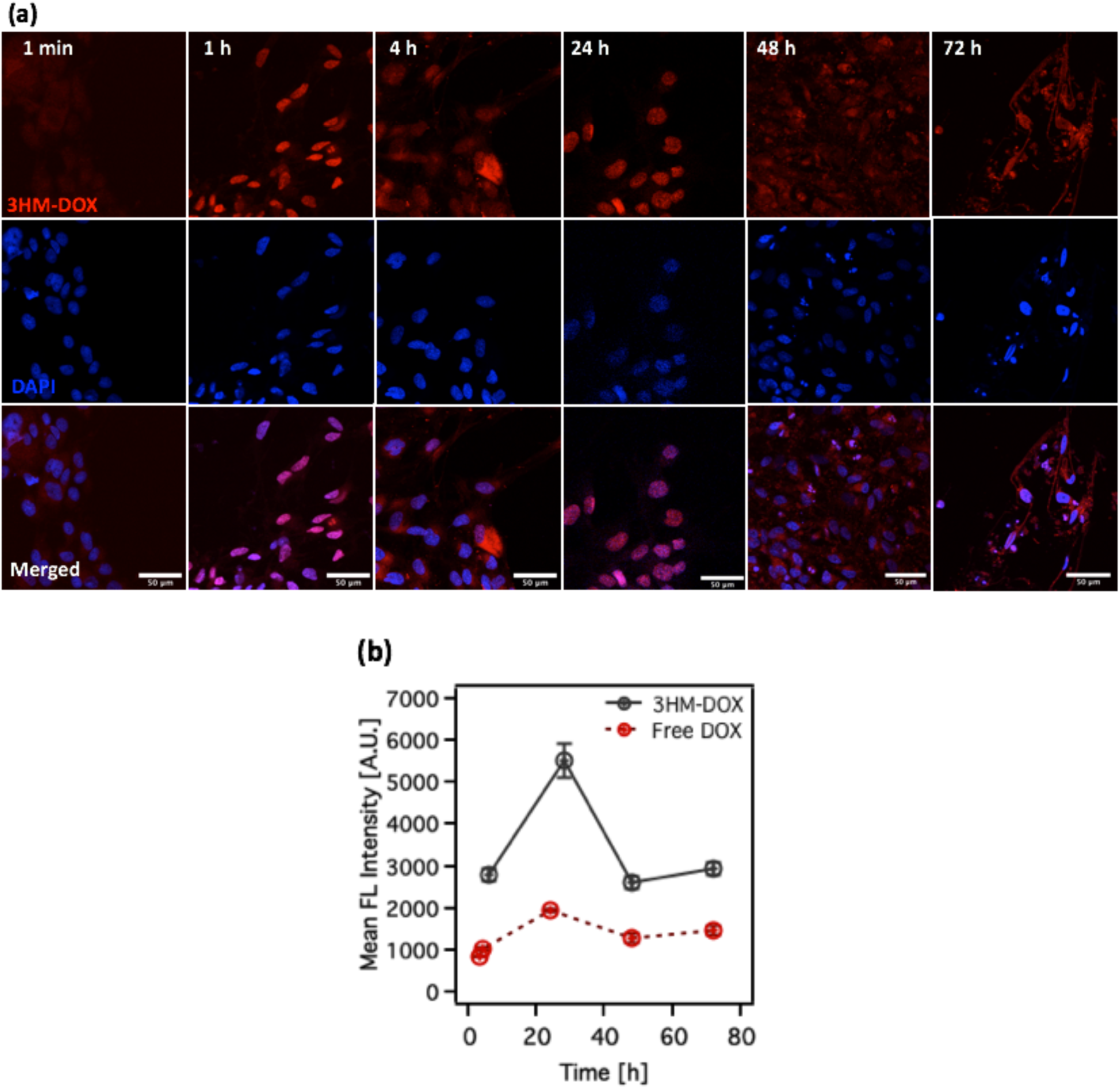
(a) Confocal fluorescence micrographs showing intracellular accumulation of DOX FL in GBM6 cells over 72 h of treatment with 3HM-DOX. scale bars are 50 µm. (b) Temporal dependence of free DOX and 3HM-DOX internalization by GBM6 cells measured by average bulk FL of DOX with FCT.

### 2.9. Determination of Internalization Pathway

3HM-FAM treatments were prepared as detailed above. GBM6 cells were seeded and incubated for 48 h to allow attachment. ^20^ min before adding treatments, cells were exposed to several small molecule inhibitors of specific cell internalization pathways (Table 1) and receptors, or reduced temperature (4 °C). This selective reduction of chosen cellular processes allow the elucidation of the major relevant pathways for 3HM internalization. ^24–26^ Appropriate inhibitor concentrations (shown in Table 1) were determined by halving the concentration when cytotoxicity became morphologically noticeable. Treatments were continuously incubated with cells for 3 h. Cells were prepared into FCT solutions as detailed above. An LSR II Analyzer (BD Biosciences) was used to measure 10,000 cells for each well. FAM fluorescence was measured at 530 nm (FITC channel) with an excitation wavelength of 488 nm. Average bulk fluorescence from each gated sample was compared to the control of uninhibited cell uptake.

**Table 1.** Concentrations of uptake inhibitors used to determine the internalization pathway of 3HM in Figure 5. ^24–26,38^

| **Uptake Pathway** | **Inhibitor (Mechanism)** | **Conc. I**<br>**[μg/mL]** | **Conc. II**<br>**[μg/mL]** |
| --- | --- | --- | --- |
| Clathrin-mediated Endocytosis | Chlorpromazine<br>(inhibit pit formation) | 5 | 10 |
| Macropinocytosis | Nocodazole<br>(microtubule disruption) | 3.75 | 7.5 |
| Macropinocytosis/Phagocytosis | Cytochalasin D<br>(inhibit actin polymerization) | 5 | 10 |
| Clathrin-mediated/ non-associated endocytosis | Monensin<br>(inhibit lysosome formation) | 0.035 | 0.07 |
| Caveolae-mediated Endocytosis | Genistein (local actin disruption,<br>dynamin II inhibition) | 12.5 | 25 |

### 2.10. Statistical Analysis

Data presented is from independent triplicate experiments unless otherwise noted. Error bars are shown as standard deviation. Welch’s t-tests (two samples with unequal variances) were performed for statistical comparisons. P values less than. 05 were considered statistically significant.

## 3. Results and Discussion

### 3.1. 3HM-DOX cytotoxicity *in vitro*

To enhance relevance of presented *in vitro* results to future *in vivo* translation, appropriate treatment concentrations and exposure times were estimated from previous 3HM biodistribution studies. ^7,10^ Accordingly, *in vivo* exposure and clearance processes were imitated *in vitro* with culture conditions consisting of a 48 h treatment period of 3HM-DOX, followed by removal of treatments, washing, replacing with fresh media, and culturing for 72 h before assaying viability. Quantifying cytotoxicity over these time scales was done with MTT metabolic assay. Figure 1A shows that 72 h after removal of treatments, 3HM-DOX demonstrated dose dependent cytotoxicity to PDX GBM6 cells, with an inhibitory concentration (IC_50_) of 6.2 µg/mL. This concentration is ~13 times higher than that of the free drug (0.47 µg/mL) but still less than predicted *in vivo* exposures. This 3HM-DOX treatment regimen was also effective on two immortalized GBM cell lines, U87MG (IC_50_ = 15.0 µg/mL) and LN229 (IC_50_ = 21.5 µg/mL), as shown in Figure 1B and 1C. U87MG showed more gradual dose dependence than LN229, more similar to GBM6. This difference in behavior puts emphasis on careful choice and characterization of cell models to ensure comparable behavior across other cell and animal studies. Treatments of matched concentrations of free DOX were more cytotoxic for U87MG (IC_50_ = 6.5 µg/mL) and LN229 (IC_50_ = 0.51 µg/mL).

The differences in cytotoxicity of free DOX and 3HM-DOX suggest that they have distinct internalization / intracellular trafficking mechanisms. DOX dose dependence correlates with nuclear availability of active DOX, based on its mechanism of action. ^27^ Free DOX can passively enter cells and traffic to the nucleus, in addition to active influx and efflux mechanisms of trafficking. ^20,27^ Nanocarriers, including 3HM (discussed later in this work), have been reported to require active uptake and trafficking processes. ^28–31^ Additional intracellular disassembly / release processes are needed before 3HM releases its cargo, allowing it to reach the nucleus. This constitutes a reduction in cell response over a comparable direct exposure.

Quantifying the cytotoxicity of 3HM-DOX formulations under direct exposure allowed the calculation of mathematically-guided *in vivo* dosing from existing biodistribution data with our recently published framework of diffusive flux modeling. ^23^ This model incorporates both plasma pharmacokinetics and tumor/ nanoparticle permeability into its analysis. Biodistribution data published with 3HM and U87MG tumor models allowed these factors to be established and suggest IC_50_ exposures found here are achievable *in vivo*. ^^7,10,12^^ Theoretical minimum single-injection doses capable of reaching IC_50_ exposures (15.0 µg/mL over 48 h) were predicted to be ~10 and ~13 mg/kg in rat and mouse models, respectively, as shown in Supporting Information Figure 3. This sort of accessible analysis helps to establish a baseline for dosing regimens in ongoing *in vivo* studies with minimal resources.

**Figure 3.**
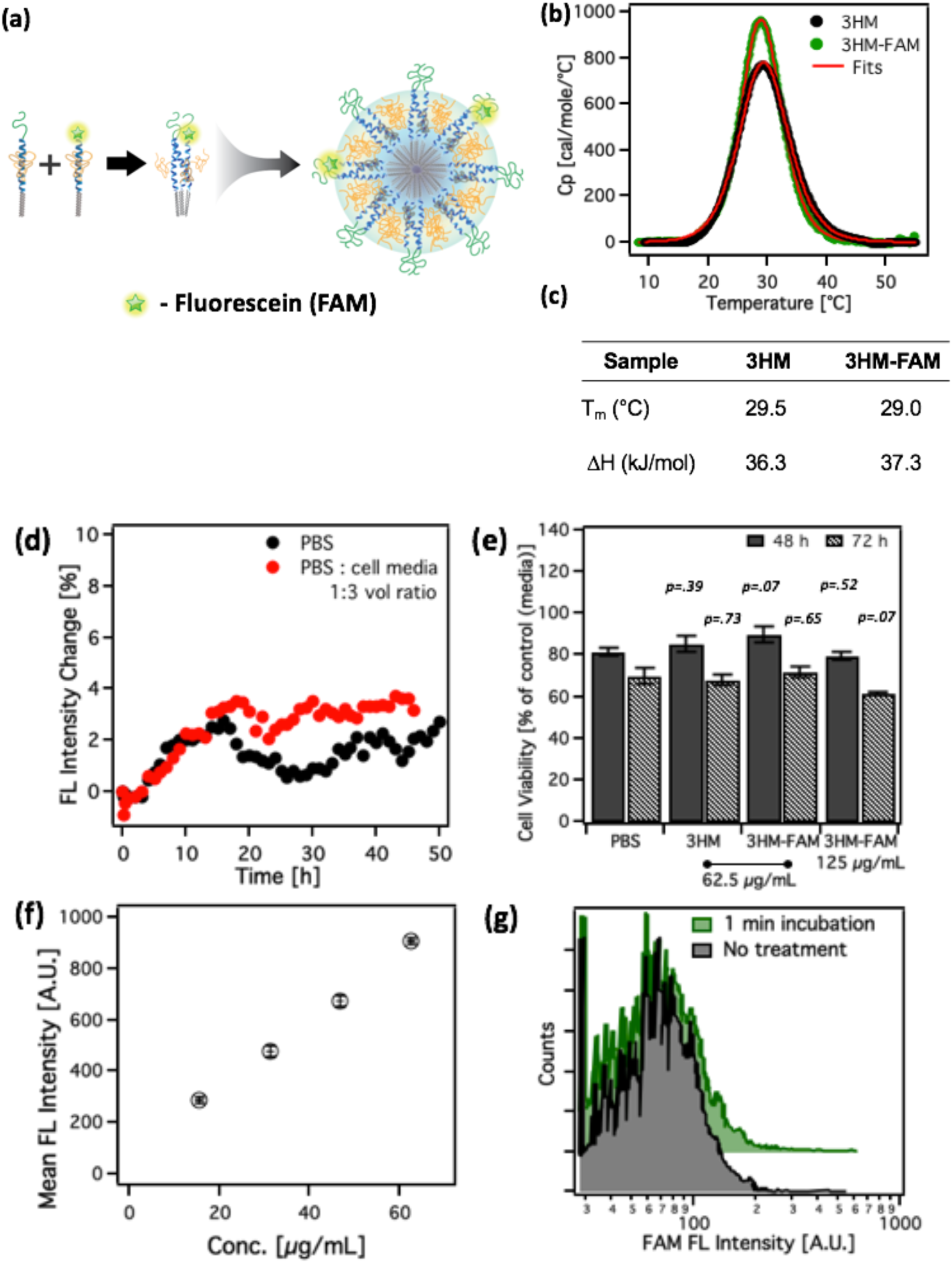
(a) Schematic of 3HM-FAM system with ~2 FL-labeled amphiphiles per micelle (4% FL-labeled). (b) Concentration normalized DSC thermograms for 3HM and 3HM-FAM. (c) Alkyl transition temperatures and enthalpies of 3HM and 3HM-FAM (d) Fluorescent recovery of 3HM-FAM in phosphate buffer and cell media over 48 h. (e) GBM6 viability with 3HM and 3HM-FAM treatment measured by MTT assay. Amphiphile concentrations are matched to the amphiphile content of 5 and 10 µg/mL 3HM-DOX formulations (8 wt%). * *p* <. 05, ** *p* <. 001 for series vs. respective PBS controls. (f) Concentration dependence of 3HM-FAM internalization by GBM6 cells. (g) Histograms of FAM signal measured by FCT of untreated GBM6 cells and cells treated with 3HM-FAM for 1 min, then subsequently washed.

### 3.2. 3HM-DOX internalization by GBM6 cells

Confocal images of GBM6 cells incubated with 3HM-DOX in Figure 2A showed DOX fluorescence localized to cell nuclei within 1 h. DOX signal inside the cells continued to accumulate to 24 and 72 h. Cytotoxic effects were also evidenced by abnormal morphologies, binucleation, and cellular debris at longer time points. The cellular uptake of 3HM-DOX was investigated over larger populations of cells with FCT. GBM6 cells treated with 3HM-DOX showed a higher signal for internalized DOX than cells treated with the free drug of the same concentration as shown in Figure 2B. 3HM-DOX, as well as free DOX fluorescence, reached a maximum at 24 h, and then decreased, matching the time scale of DOX consumption as it goes through its cytotoxic action. Encapsulation of DOX into 3HM enhanced its uptake, despite free DOX showing greater cytotoxicity than 3HM-DOX above in Figure 1. These two observations indicate intact 3HM-DOX particles were uptaken and retained cargo intracellularly; DOX content is higher inside cells treated with 3HM-DOX, but not transporting to the nucleus or going through its cytotoxic action as quickly as free DOX. This also suggests 3HM-DOX is not liberating DOX extracellularly or immediately upon internalization. 3HM formulations must be processed through intracellular machinery distinct from free drug, before its cargo becomes available. The conditions, timescales and energy scales of 3HM-DOX disassembly and DOX release are currently under investigation for future publication.

### 3.3. 3HM-FAM characterization and cellular internalization

After confirming the internalization and efficacy of 3HM-DOX treatments, the uptake of 3HM amphiphiles themselves was investigated. Analysis is convoluted if relying on DOX fluorescence to track 3HM-DOX due to DOX elimination as it goes through its mechanism of action (Figure 1B), and physical encapsulation inside of 3HM. To explicitly understand the uptake of 3HM itself, 5,6-carboxyfluorescein (FAM) was covalently attached to the 3HM amphiphile. To ensure that fluorescently-labeled 3HM showed representative behavior of the unlabeled construct, FAM labelled amphiphiles were doped into blank 3HM amphiphiles for cell uptake studies at 4 mol% (~2 FAM per micelle), yielding fluorescent micelles (3HM-FAM) as shown in Figure 3A. This minimized any change to micelle stability or surface properties, since FAM replaced the surface PEG in the amphiphile architecture. The alkyl core melting transitions of 3HM and 3HM-FAM were comparable in Figure 3B and 3C, displaying similar melting temperatures of 29.5 and 29.0 °C, and enthalpies of transition of 36.3 and 37.3 kJ/mole/°C respectively. Size, as measured by dynamic light scattering (DLS), was unchanged as a result of FAM-labeled amphiphile incorporation as shown in Supporting Information, Figure 4. The kinetic stability of 3HM-FAM was probed with fluorescence of bulk solutions in phosphate buffer and in 1:3 phosphate buffer:cell media (cell treatment ratio). Figure 3D shows similar nominal increase in fluorescence over 48 h in both conditions, confirming 3HM-FAM was fluorescently stable and free from quenching or release effects over experimental timescales. 3HM-FAM did not show cytotoxicity against GBM6 cells at concentrations of amphiphile matched to the amphiphile content of 5 and 10 µg/mL 3HM-DOX formulations, compared to unlabeled 3HM and PBS, as shown in Figure 3E. This indicated the cells had similar viability and metabolism when treated with 3HM-FAM as with blank 3HM.

**Figure 4.**
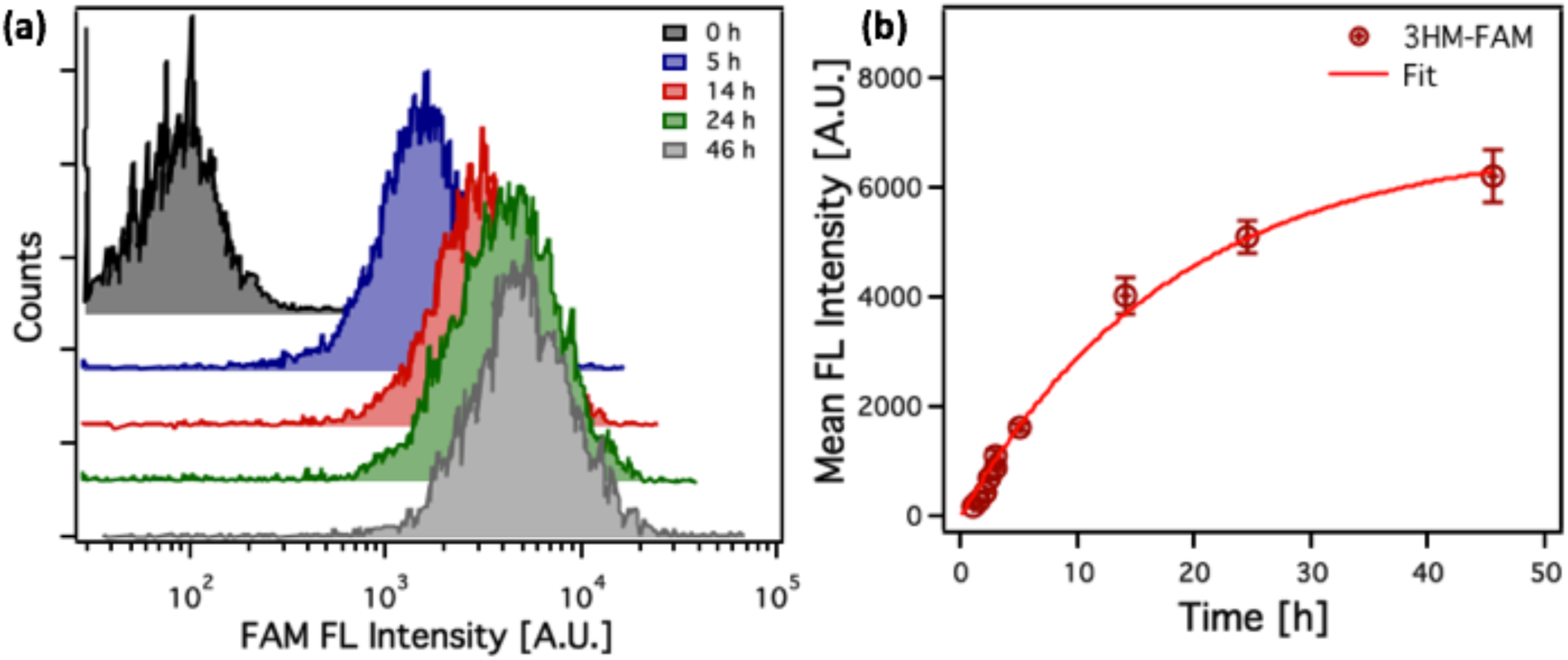
(a) Representative histograms of FAM FL of GBM cells treated with 3HM-FAM (4% FL-labeled) over time measured by FCT. (b) Temporal dependence of 3HM-FAM (4% FL-labeled) internalization by GBM6 cells showing internalization t_1/2_ = 12.6 h measured by average bulk FL of FAM with FCT.

Dose dependence of cell internalization of 3HM-FAM in Figure 3F was observed over amphiphile concentrations matched to previous 3HM-DOX experiments. Since FAM on its own cannot enter cells (Supporting Information, Figure 5), it is certain that fluorescence signal is from intact FAM-labelled amphiphiles. ^32^ Controlling for the effects of surface-adsorbed micelles was done by comparing untreated GBM6 cells with 3HM-FAM treated cells for 1 minute, followed by PBS washing before preparing the cells into FCT sample solutions. Figure 3G shows the insignificant differences between these treatments, verifying that fluorescence signal was a result only of internalized amphiphiles.

**Figure 5.**
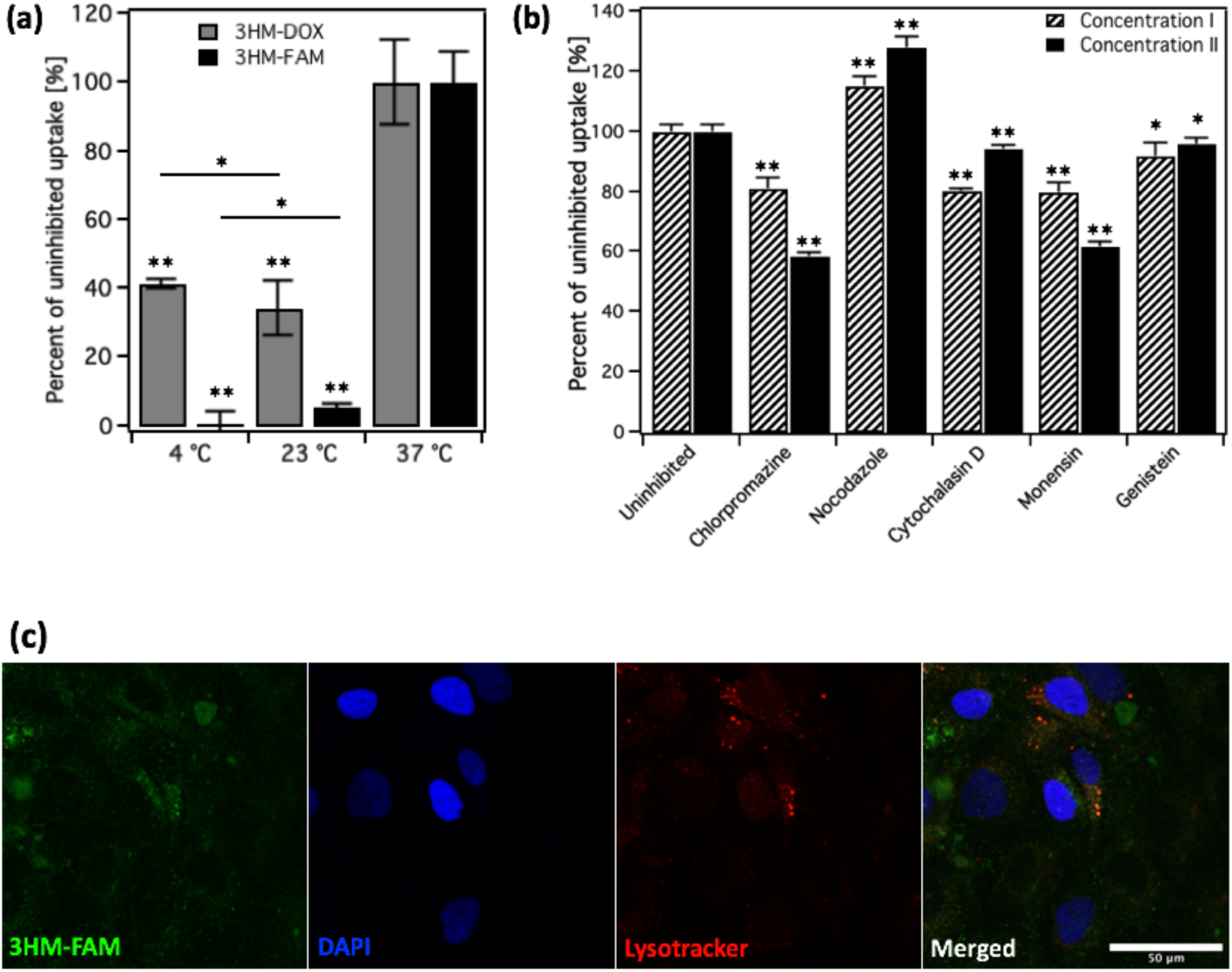
(a) 3HM-FAM (4% FL-labeled) and 3HM-DOX internalization by GBM6 cells depending on temperature. Uptake is normalized to 100% for 37°C. (b) 3HM-FAM co-incubation with indicated molecular inhibitors of various uptake pathways, as measured by average bulk FL of FAM with FCT. Uptake is normalized to 100% for respective uninhibited controls. (c) Confocal fluorescence micrographs showing colocalization of 3HM-FAM nanocarriers with lysosomes after 24 h incubation. Scale bar is 50 µm. For statistical comparison: * *p* <. 05, ** *p* <. 001 for series vs. ^37^ °C control, unless denoted in (a), and for series vs. respective uninhibited controls in (b).

Cellular uptake of 3HM-FAM over 48 h showed population-wide uptake of 3HM-FAM in Figure 4A. No cells were observed to retain low fluorescent signal over 48 h. Low broadening of the distribution of fluorescent intensity with 3HM-FAM internalization also indicates homogenous uptake by the majority of cells. Figure 4B displays the tracking of the mean fluorescence intensity of these GBM6 cells over time. Fitting with one phase association kinetics resulted in a K_int_ of 0.055 (internalization t_1/2_ = 12.6 h). This internalization rate was notably slower than some other nanoparticle systems, especially given the size and self-assembled nature of 3HM. Liposomes around 100 nm showed internalization t_1/2_ of 3-6 h in F98 and U-118 MG cells. ^^33^^ Dendrimers of 150 nm showed an internalization t_1/2_ of <5 h in J774 cells. ^^34^^ Polylactic acid (PLA) particles around 100 nm showed an internalization t_1/2_ of 1.2–2.5 h in RG2 cells. ^^35^^ It is theorized cell line metabolism (doubling time) and nanoparticle physical characteristics (size, composition, surfaces properties etc.) are partially responsible for these differences, but the distinction of these kinetics from 3HM is still significant. The longer internalization time, coupled with homogeneous uptake of 3HM across different cell subpopulations of GBM6 cells, suggested a regulated active uptake process is involved.^36^

### 3.4. 3HM cellular internalization mechanism

To investigate the mechanism of internalization of 3HM, uptake was measured under reduced incubation temperature, which has been shown to retard all active cellular uptake mechanisms. ^4,24,37^ Diffusion (passive uptake) is also reduced but has been proved to not be the limiting factor in cell uptake at lowered temperature, even to 4 °C. ^24,37^ Incubation times at lower temperatures were limited to 3 h, and cells were still well-adherent after these times (Supporting Information, Figure 6). Interestingly, 3HM-FAM internalization by GBM6 cells was almost entirely shut down at both 23 °C and 4 °C (94% and 100%, respectively), indicating almost all 3HM uptake occurred by active processes, as shown in Figure 5A. 3HM-DOX uptake was also significantly reduced by lowered temperature (66% and 59%, for 23 °C and 4 °C respectively).

**Figure 6.**
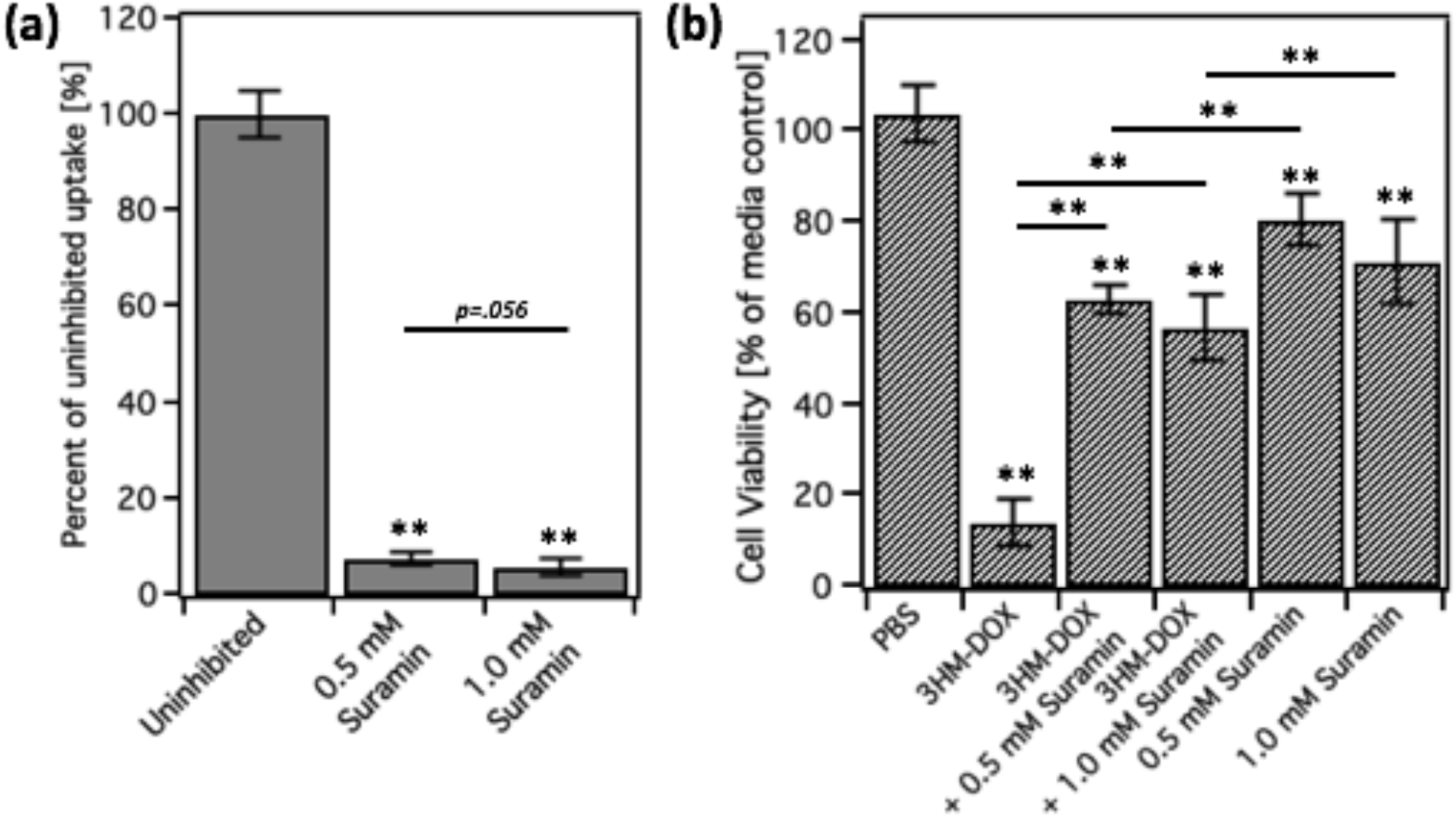
(a) FCT measurements showing 3HM-FAM (4% FL-labeled) internalization by GBM6 cells is inhibited when LDLR is blocked with suramin over 3 h incubation. * *p* <. 05, ** *p* <. 001 for series vs. uninhibited control unless denoted. (b) Viability (MTT assay) of GBM6 cells treated with 3HM-DOX (10 µg/mL DOX, 48 h treatment, 72 h post incubation) and / or suramin to confirm 3HM uptake inhibition. * *p* <. 05, ** *p* <. 001 for series vs. PBS control unless denoted.

The specific active uptake pathways involved in 3HM uptake were examined by co-incubation with chemical inhibitors. Since these inhibitors can alter other cell functions besides uptake phenomenon, cell exposure was limited to conservative concentrations, as reported in Table 1. ^24,25,38^ Two concentrations of inhibitors were used based on viability, with the highest concentration being half of the concentration where cell morphology began to change (images shown in Supporting Information, Figure 6). ^24,25^ Cell uptake of 3HM-FAM after a relatively short 3 h of co-incubation is shown in Figure 5B. Inhibiting clathrin-dependent and non-associated endocytosis reduced 3HM uptake the most, indicating these are the most prominent pathways in its internalization. Total inhibition was not observed (or desired) for any particular pathway in these experiments due to the modest amounts of inhibitors used. Lysotracker staining showed colocalization of 3HM-FAM with lysosomes (downstream of endosomes) in GBM6 cells as shown in Figure 5C, confirming an active uptake mechanism. The involvement of these endocytosis pathways is interesting since they are usually receptor-initiated pathways, and 3HM has no designed specific chemical moieties. Protein corona formation likely contributes to the ability of 3HM to enter through these often receptor mediated pathways. ^4,31,39,40^ Additionally, the absence of (macro)pinocytosis and phagocytosis involvement in uptake for GBM6 may contribute to favorable 3HM performance *in vivo*, since this is the major mechanism of how macrophages and other phagocytes remove particle from circulation. These pathways are not active in internalizing 3HM, so reduced removal by phagocytes likely allows for the longer circulation observed in animal models. ^4,24^ Similar reduction in 3HM uptake with clathrin-mediated endocytosis inhibition was observed with U87MG cells (Supporting Information, Figure 7A).

### 3.5. Involvement of receptor surface recognition in 3HM internalization

3HM’s PEGylated surface is predicted to attract a similar protein corona composition to other PEGylated particles based on comparison of its surface characteristics. ^29,41,42^ The density of brush layer PEG750 on 3HM is ~7 chains per 100 nm^2^ based on geometric calculations with previously determined internal structural information of the trimeric peptide headgroup and PEG2K conformation. ^13,15^ If only considering these chains, the density is below the 20 chains per 100 nm^2^ reported as a minimum threshold for circulation prolongation. ^39^ However, the areas between the PEG750 presented by the peptide is also composed of PEG2K compressed out from the side-conjugated position, which brings it above this threshold. ^13,14,39^ The resulting surface thus has two different PEG conformations shown in Scheme 1, (1) lateral correlation of the PEG750, which corresponds to the interhelical distance between α-helical headgroups, either within a trimeric coiled-coil or between trimers (1.2 and 3.5 nm respectively), 15 and (2) a hydrated PEG2K surface filling in the space between trimers. ^10,13,15^

PEGylated surfaces (even below this 20 chains per 100 nm^2^ threshold) have been reported to attract elevated levels of apolipoproteins to their protein coronas, notably apolipoprotein B-100 (ApoB-100) and apolipoprotein E (ApoE). ^29,40–42^ These two moieties can be recognized by the LDLR (low density lipoprotein receptor), which is integral to the internalization of low density lipoprotein (LDL) particles. ^43^ LDL particles are self-assembled ^18–25^ nm particles (comparable size to 3HM) that contain one copy of ApoB-100 (65 nm long, pentapartite, and flexible), which envelops and stabilizes a lipid core. ^43^ The regularly patterned PEG surface of 3HM could potentially allow ApoB-100 (or other apolipoproteins) to surround it in serum-containing media, as it does in its state with similarly-sized natural LDL particles. As a self-assembled system with similar size and physical characteristics to LDL particles, this apolipoprotein-coated 3HM could behave like a synthetic LDL, at least at the cellular transport level. ^29,40,41^ Ongoing studies aim to assess PEG conformation and topology at the surface of 3HM and identify the protein corona moieties on 3HM in a quantitative manner to support this mechanism of uptake.

To investigate the involvement of LDLR in 3HM internalization, uptake and cell viability was investigated with LDLR inhibition, by the means of a known chemical inhibitor, suramin. ^44,45^ Inhibiting LDLR with modest exposures (10X less that reported ^44,45^) of suramin reduced 3HM-FAM uptake by 93% and 94% (for 0.5 mM and 1.0 mM suramin, respectively) in GBM6 cells as shown in Figure 6A. This phenomenon was also confirmed in U87MG cells with 92% uptake reduction by 1.0 mM suramin (Supporting Information, Figure 7B). This showed that surprisingly, 3HM is heavily reliant on LDLR to initiate internalization, despite not having any designed surface targeting. Importantly, since suramin inhibition happens extracellularly, off-target cellular machinery is less-affected than inhibitors used in Table 1. Since the FAM conjugation site is close to the surface, and FAM could potentially alter specific surface interactions, 3HM-DOX cytotoxicity was also studied with LDLR inhibition to confirm recognition of 3HM with a fully PEGylated surface. Figure 6B shows suramin (0.5 mM and 1.0 mM) was able to recover cell viability by 55% and 48%, respectively, when co-incubated with 3HM-DOX (10 µg/mL DOX, above 1C_50_) with the same 48 h dose - 72 h recovery regimen in previous experiments. This recovery is even more pronounced when compared to the controls of treatment only with suramin (74% and 75%, for 0.5 mM and 1.0 mM respectively). This proved that with the viability effect of suramin controlled (which could be involved in the 3HM-FAM cell uptake experiments), 3HM-DOX uptake was blocked through LDLR inhibition.

Systems derived directly from natural LDL components have been reported and manipulated for therapeutic use. ^40,44,45^ While 3HM contains similar components (peptide and alkyl chains), it does not include any of the comprising molecules of LDL, especially ApoB-100, or any features specifically recognizable by LDLR. Procurement of a nanomaterial with appropriate physical characteristics is enough to elicit specific recognition and manipulation, once populated with a preferential protein corona. Laterally-correlated surfaces could be used to induce varying biological responses, if design rules for proper biasing of bound proteins can be developed. The design and thorough characterization of nanomedicine systems that transport actively are becoming more imperative. Researchers are showing that active cell processes are more responsible for nanomaterial trafficking at cell tissue and system levels than previously thought, even for PEGylated particles that were presumed to be passively-transported. ^40,41,46^ The implications of transporting 3HM through natural machinery is currently being studied at the transcriptional level in order to provide insights on alterations that might not be clear when looking just at therapeutic outcomes, such as viability or uptake. In the context of GBM, LDLR is reported to be upregulated in many GBM tumors (128,000 – 950,000 receptors/cell) and negligible in healthy brain cells. ^44,47^ Further studies are warranted to determine if 3HM formulations can be preferentially internalized by GBM cells once transported to the brain tissue.

### 4. Conclusions

3HM-DOX was found to enhance the accumulation of DOX intracellularly in PDX GBM6 cells, which is advantageous for longer term and *in vivo* experiments where all of intracellularly accumulated DOX has more time to release and transport to the nucleus. 3HM-DOX formulations displayed dose dependent cytotoxicity to GBM6 (IC_50_ = 6.2 µg/mL), U87MG (IC_50_ = 15.0 µg/mL), and LN229 (IC_50_ = 21.5 µg/mL) cells. Published and concurrent *in vivo* studies indicate these concentrations are achievable using long-circulating 3HM in rodent models with intravenous delivery. 3HM was found to accumulate in GBM6 cells with a half-life of 12.6 h, relatively long considering its size and self-assembled design. 3HM nanocarrier formulations were found to be endocytosed uniformly by GBM6 cells, mainly via clathrin-mediated pathways, despite having no targeting moieties. LDLR inhibition by suramin was found to reduce 3HM uptake by 93 % in GBM6 cells, suggesting that 3HM is able to traffic through autologous LDL particle machinery. These findings emphasize the need for nanocarriers to be stable over long times, since cell uptake can take hours to accumulate in abundance, without even considering upstream transport stages that must be endured *in vivo*. The present findings contribute to ongoing efforts to elucidate and control the ultimate fate of 3HM in the body and to pursue informed dosing regimens based on knowing the temporal distribution of 3HM at the cell, tissue, and system levels. They also establish a standard of biological performance for the current design architecture of 3HM. Future designs of 3HM variants will take advantage of these insights to engineer desired protein corona-3HM interactions and thus subsequent protein corona-cell interactions. Results presented here are encouraging for the use of 3HM as a controlled delivery platform to GBM and potentially to other tumors.

## ASSOCIATED CONTENT

### Supporting Information

This material is available free of charge via the Internet.

MALDI-TOF, DFM analysis, Size by DLS, FCT controls, and U87MG cell uptake data (PDF).

## Author Contributions

B.T.J. and T.X. formulated the project. B.T.J., K.J., M. Lim, and M. Li synthesized and characterized materials used. R.S. and T.O. provided tissues and training for cell experiments.

B.T.J. and K.J. performed cell experiments. T.X. guided project progress. The manuscript was written through contributions of all authors. All authors have given approval to the final version of the manuscript.

## Notes

The authors declare no competing financial interest.

## Acknowledgements

Flow Cytometry was performed at the Flow Cytometry Facility of the Cancer Research Laboratory (CRL) at UC Berkeley. Confocal imaging experiments were conducted at the CRL Molecular Imaging Center, supported by NSF (DBI-1041078). PDX flank tumors were obtained from the Preclinical Therapeutic Testing Core Facility at the Brain Tumor Center at UC San Francisco.

