## Supporting Information and Figures for "Design of 18 nm Doxorubicin-loaded 3-helix Micelles: Cellular Uptake and Cytotoxicity in Patient-derived GBM6 Cells"

**Number of pages: 7**

**Number of figures: 7**

**Number of tables**: **0**

*Supporting Methods*

**Matrix-Assisted Laser Desorption Ionization.**

3 μL of sample (1 mg/mL) was added to an equal volume of α-cyano-4-hydroxycinnamic acid (10 mg/mL) matrix solution (50:50 $H_2O$:ACN), the mixture was spotted and dried for 30 minutes on a stainless steel MALDI plate and the spectrum was collected on a MALDI-TOF (Applied Biosystems).

**Diffusive Flux Modeling Analysis**

To predict the minimum *in vivo* single-injection intravenous dose required for eliciting tumor toxicity based on *in vitro* $IC_{50}$ values, 3HM-DOX tumor accumulation curves were projected using our published diffusive flux modeling (DFM) method. [1] For intracranial U87-MG xenografts, effective permeability of 3HM was found to be 0.670 μm/h. This value was used in the DFM equation along with 3HM pharmacokinetic parameters in rodents (i.e. $C_v = C_0 \exp(-0.0454t)$ for rats, and $C_v = C_0 (0.281 \exp(-1.508t) + 0.719 \exp(-0.024t))$ for mice). $C_v$ is the 3HM-DOX blood vessel concentration (μg/μL) which changes over time, while $C_0$ is the initial concentration (μg/μL), and t is time (h). Fraction of available volume in tumor interstitial is 0.173 for 3HM in rats, and 0.149 for mice. Weights are assumed to be 294 g for rats and 25 g for mice, with initial blood concentration of 6.5% injected dose per $cm^3$ (%ID/cc) in rats and 63 % ID/cc in mice. In this way, theoretical doses administered in mg/kg could be translated into $C_0$ in μg/μL.

**Dynamic Light Scattering.**

Size distributions of micelles with and without 3HM-FAM doping were measured using a BI-200SM light scattering system (Brookhaven Instruments; laser: 637 nm, 30 mW; 90° scattering geometry, 100 μm filter, 25 °C). Three autocorrelation functions of the scattered light were collected for 1 minute each and analyzed to determine number average size distribution.

**Optical Microscopy.**

Optical microscopy was performed with an Olympus BX51 microscope. GBM6 cells were seeded on 12-well plates and incubated for 48 h. Cells were incubated with chemical inhibitors of the concentrations indicated in Table 1 for 3 h. Wells were imaged to look for morphological signs of effects on off-target cellular processes.

*Supporting Figures*

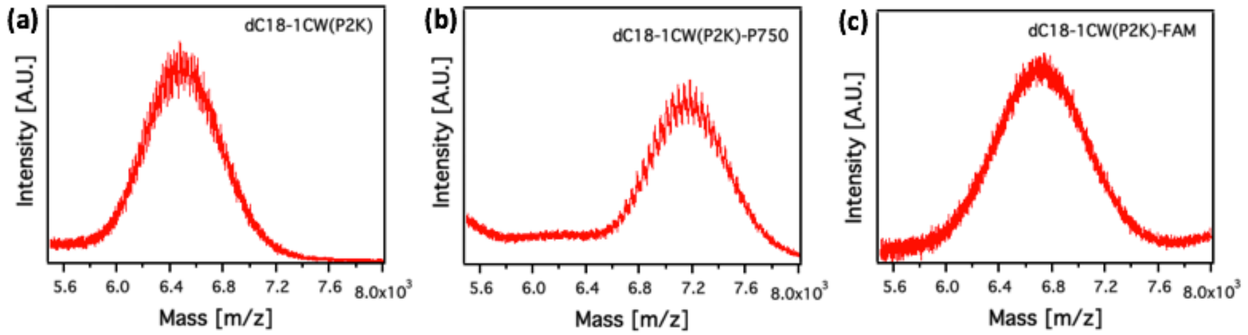


**SI Figure 1:** MALDI-TOF mass/charge spectrums of (a) dC18-1CW(PEG2K) [max: 6474 g/mol], (b) dC18-1CW(PEG2K)-P750 [max: 7178 g/mol, 44 g/mol spacing between P750 monomers], and (c) dC18-1CW(PEG2K)-FAM [max: 6734 g/mol].

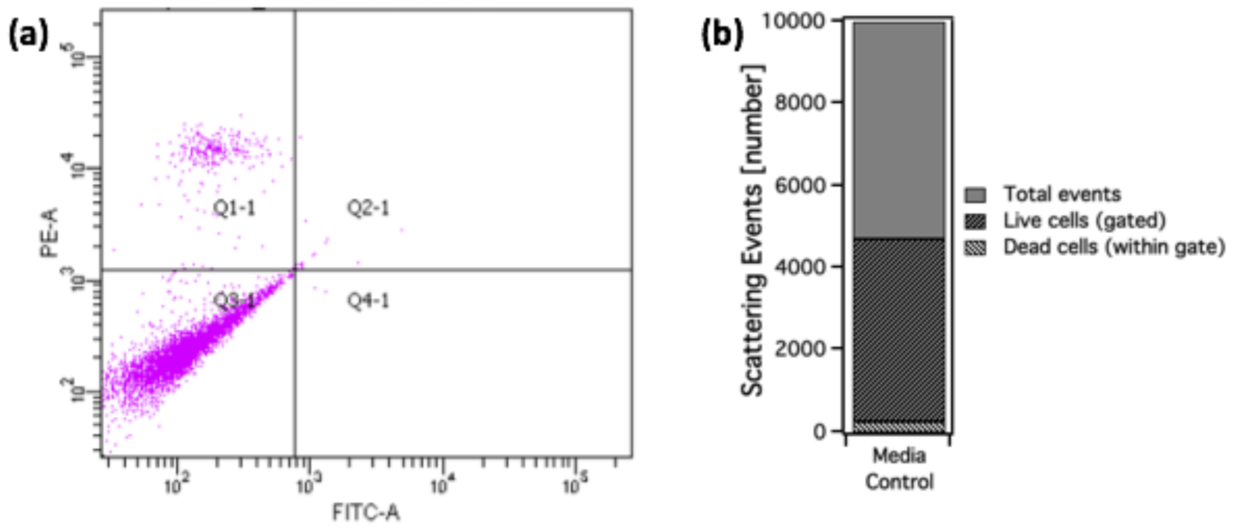


**SI Figure 2:** Flow cytometry gating validation with propidium iodide (PI) staining. (a) scatter plot of untreated GBM6 cells co-incubated with PI for 3 h. PI fluorescence (measured in PE channel) indicates non-viable cells. (b) breakdown of gating for all events in one measurement.

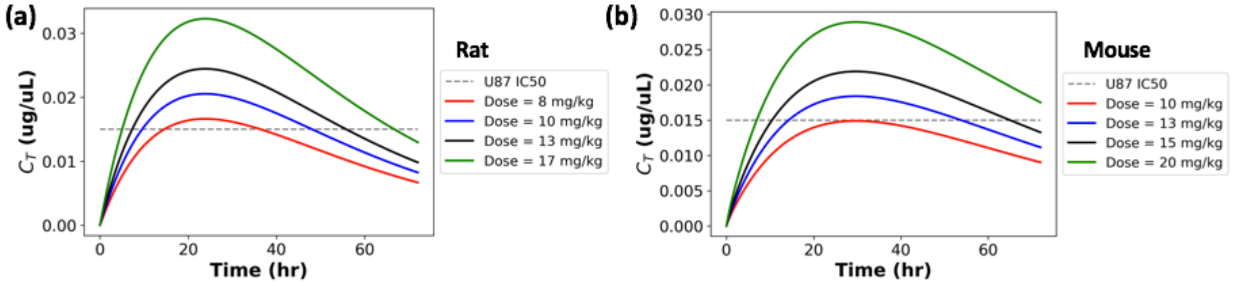


**SI Figure 3:** Theoretical projected tumor concentration profiles of *in vivo* single-injection dosing regimens based on diffusive flux modeling analysis applied to (a) rat and (b) mouse models bearing U87MG tumors. Critical parameters: U87MG $IC_{50}$ = 15.0 μg/mL; effective permeability ($P_{eff}$) = 0.67 μm/h (all other modeling parameters as specified in Reference [1]).

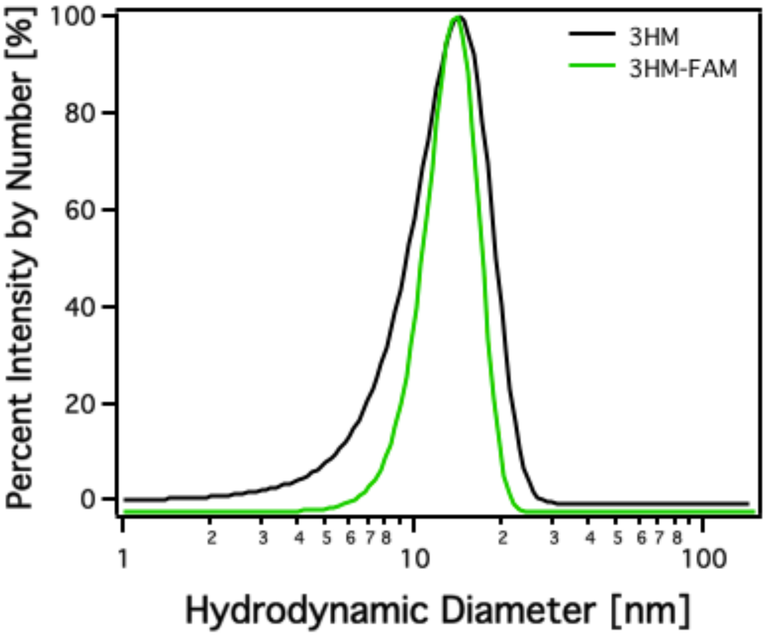


**SI Figure 4:** Size distributions of micelles with and without 3HM-FAM doping measured by DLS. Number average diameter is 17 nm.

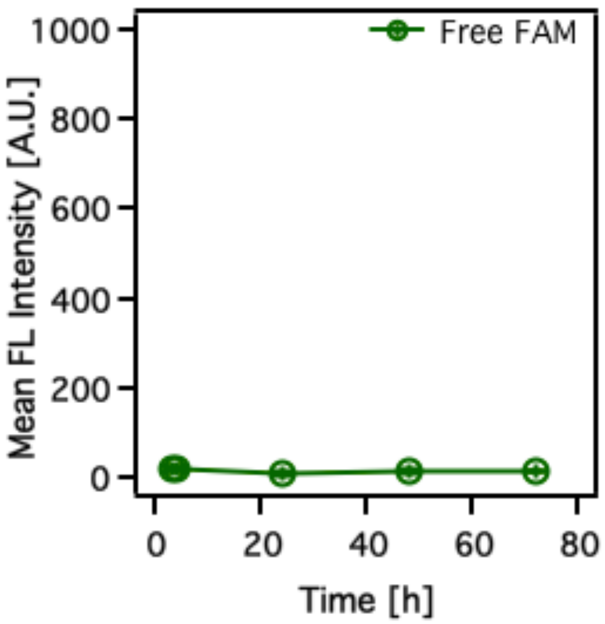


**SI Figure 5:** Temporal dependence of free FAM internalization by GBM6 cells showing no cell uptake by average bulk FL of FAM with flow cytometry.

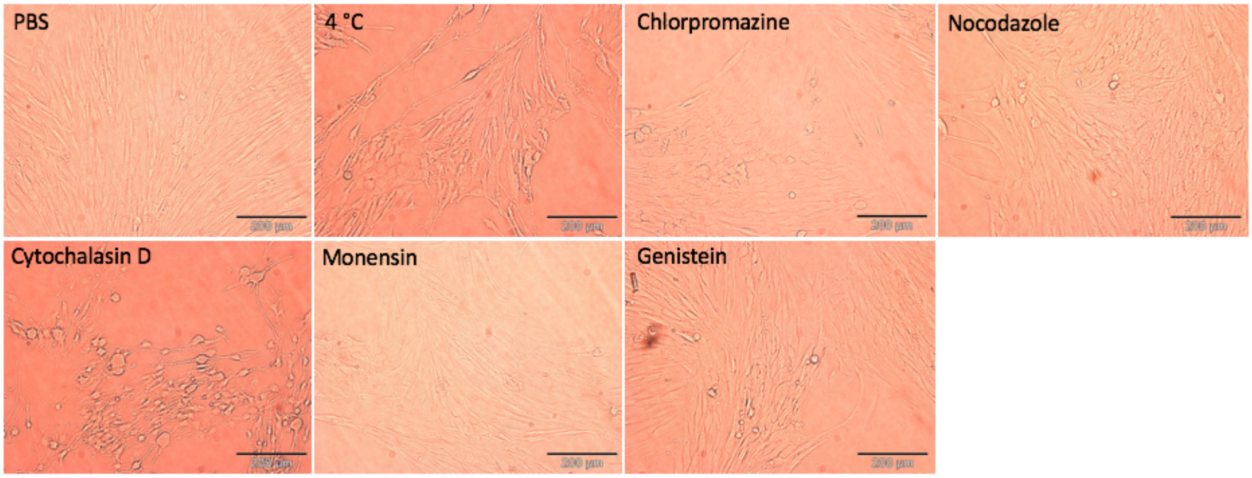


**SI Figure 6:** Optical micrographs showing GBM6 cell morphology under indicated inhibitory conditions after 3 h incubation. Concentrations used for coincubation are the 'Concentration II' indicated in Table 1 in the main text. Scale bar is 200 µm.

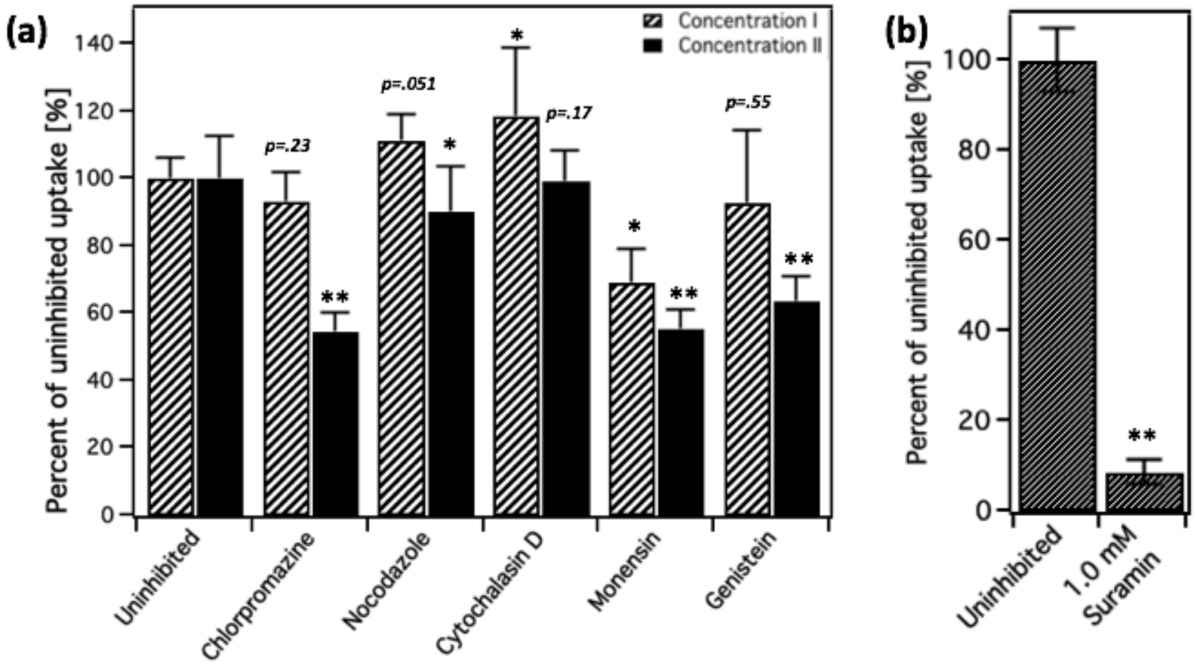


**SI Figure 7:** 3HM-FAM (4% FL-labelled) internalization by U87MG cells depending on (a) co-incubation with indicated molecular inhibitors of various uptake pathways measured by average bulk FL of FAM with flow cytometry (Concentrations are listed in Table 1 in the main text) and (b) co-incubation with suramin, an inhibitor of LDLR. For statistical comparison: * $p < .05$, ** $p < .001$ for series vs. uninhibited uptake.

***Supporting Information References***

1. Lim, M.; Dharmaraj, V.; Gong, B.; Jung, B. T.; Xu, T. Estimating Tumor Vascular Permeability of Nanoparticles Using an Accessible Diffusive Flux Model. *ACS Biomaterials Science & Engineering* **2020**, DOI: 10.1021/acsbiomaterials.9b01590.
